# Dual-reporter assay for selective small molecule inhibition of expanded toxic huntingtin

**DOI:** 10.64898/2026.11.06.764921

**Authors:** Elizabeth A. Kraichely, Sungah Kim, Jen-Chieh Wang, Kyung Woon Jung, Peter J. Chung

**Author notes:** **Author Contributions:** E.K., K.J. and P.C. designed experiments. E.K., S.K., and J.W. performed experiments. All authors consulted together on the interpretation of results and preparation of the manuscript. **Competing Interest Statement:** The authors declare no competing interest.

## Abstract

Huntington ‘s disease (HD) is a neurodegenerative disorder caused by an expansion of CAG repeats in exon 1of the human huntingtin gene (*HTT*). The resultant pathogenic allele (*mHTT*) produces a mutant huntingtin protein (mHtt) containing an expanded polyglutamine tract. Because mHtt drives neurodegeneration while wildtype huntingtin (Htt) remains critical for regulation of gene transcription and cell survival, an ideal therapeutic would specifically downregulate mHtt expression while preserving Htt. Small molecules are attractive candidates for HD given their ability to penetrate the blood-brain barrier and amenability to oral dosing, but there remains no high-throughput platform that reports both efficacy and allele selectivity. Here, we present a bacterial assay in which exon 1 of *HTT* and *mHTT*, each fused to spectrally distinct fluorescent proteins, are co-expressed in the same cell, allowing simultaneous quantification of their expression in the presence of small molecules. This high-throughput assay isolates the effect of small molecules on huntingtin expression without confounding factors present in mammalian cells, such as endogenous Htt or human RNA-binding proteins. To validate this assay, we evaluated two previously reported CAG repeat binding compounds and used circular dichroism spectroscopy and cell-free protein synthesis to investigate their underlying mechanisms. Furamidine exhibited reproducible, sequence-dependent effects, altering the structural stability of *HTT* RNA and selectively decreasing the wildtype Htt expression (a selectivity profile opposite to what is therapeutically required). In contrast, myricetin did not affect *HTT* RNA stability and its decreased signal in the dual-expression assay was due to fluorophore artifacts. This highlights the importance of complementary workflows in distinguishing genuine modulators from reporter-specific artifacts. Collectively, these findings establish the dual-expression assay as a screening platform for CAG repeat-targeting small molecules and demonstrate the importance of determining which allele reported compounds preferentially affect.

## Introduction

Trinucleotide repeats are ubiquitous in the human genome and have the propensity to expand, leading to the development of numerous neurodegenerative disorders^1,2^. Of these disorders, Huntington ‘s disease (HD) is the most common, caused by an expansion of CAG repeats in exon 1of the huntingtin gene (*HTT*)^1,3,4^. Physiologically, CAG repeats in the *HTT* gene typically range from 10 to 26, but individuals that develop the disease often have 40 repeats or greater with increasing CAG repeat tract length directly correlating to earlier onset of clinical symptoms associated with HD^5–7^. Indeed, the expression of this mutant huntingtin (mHtt) has been strongly associated with toxic gain-of-function pathology in mice and human stem cell models of HD (with decreased expression of mHtt shown to mitigate pathology)^8–11^. Genetic approaches, including RNA interference and antisense oligonucleotides (ASOs), aim to lower mHtt expression by degrading or silencing the mutant mRNA transcript^12–15^. Unfortunately, due to their highly charged nature, nucleic acid therapeutics are unable to easily cross the blood-brain barrier and penetrate neuronal membranes^16,17^. Further, as these approaches are designed to target RNA sequences, allele-selectivity is difficult to achieve. This can lead to off-target reduction of the wildtype huntingtin (Htt), which is critical in transcriptional regulation and cell survival^18,19^.

Small molecule therapeutics offer a compelling alternative. They can be selected for blood-brain-barrier permeability and, critically, may exploit the more thermodynamically stable RNA hairpin structure that the *mHTT* transcript adopts, exposing the inner groove for selective *mHTT* targeting^20–26^. Two small molecules, furamidine and myricetin^27,28^, have been reported to bind to CAG repeat RNA and reduce huntingtin expression, making them natural reference compounds to evaluate in a potential screening platform. Furamidine has been shown to reduce expression of the huntingtin protein, although biophysical experiments indicate that furamidine binds more strongly to shorter CAG repeat transcripts (CAG_18_) than to longer, pathogenic repeats (CAG_41_), a selectivity profile opposite to what is therapeutically required. Myricetin was found to interact with the stem of the RNA hairpin structure to reduce expression of Htt; however, it exhibited limited selectivity for expanded over non-expanded CAG repeats. The limitations of these compounds are not incidental as they trace to the assays used to identify them. The discovery of furamidine required computationally intensive *in silico* identification to screen for CAG repeat binder candidates that were subsequently tested for binding to RNA but not for selective inhibition of specific alleles. Likewise, the discovery of myricetin was the result of 36 small molecules being screened via a fluorescence titration assay against a library of 11 RNA sequences containing a short 5 ‘C**N**G/3 ‘G**N**C motif. In both cases, binding and functional readouts were obtained separately with no measurement of allele selectivity. This fragmented approach makes it difficult to discover therapeutically relevant candidate compounds as efficacy in downregulation and selectivity is not monitored simultaneously. Therefore, it is unsurprising that the assays used in these studies found CAG repeat binders unable to preferentially inhibit expression of mHtt. To address this barrier, we developed a high-throughput bacterial dual-expression assay designed for simultaneous monitoring of wildtype and mutant Htt expression, enabling rapid, quantitative assessment of both efficacy and selectivity in a basic plate reader. By using a bacterial system, this assay not only remains accessible to most laboratories without specialized infrastructure, but also eliminates confounding variables present in mammalian systems, such as endogenous *HTT* and RNA-binding proteins (like MBNL1) that interact with CAG repeat RNA^22,29^. To validate this assay, we applied furamidine and myricetin as reference compounds and complemented our cellular assay with additional biophysical assays. These included circular dichroism and cell-free protein synthesis to assess direct interaction and biochemical mechanism of action, respectively. Together, this integrated assay and workflow provides a platform for identifying and characterizing CAG repeat-targeting small molecules prior to more complex tissue models. Therefore, this assay may be utilized as an early step toward allele-selective therapeutics for HD.

## Results

### Dual-expression assay design

We designed a dual-expression bacterial plasmid encoding exon 1 of wildtype huntingtin containing either CAG_16_ (*HTT*) or mutant huntingtin containing 41 CAG repeats (*mHTT*), each C-terminally fused to a spectrally distinct fluorescent reporter (Fig. 1a, b). Fluorescent proteins mCherry2 and mGreenLantern were chosen due to their relatively short maturation times compared to the total experiment length (22.8 min and 13.5 min, respectively), high quantum yields, and distinct excitation and emission spectra to minimize fluorophore crosstalk^30,31^ (with “swapped reporter “ control experiments to rule out fluorophore artifacts, see Fig. S1). Each gene is independently regulated by its own lac operon controlled T7 promoter, allowing for inducible expression in response to the addition of isopropyl β-D-1-thiogalactopyranoside (IPTG) to produce comparable transcript levels (Fig. S2b) to independently monitor each protein ‘s expression using its fluorophore reporter (Fig. 1a). Both genes were encoded into a pET28a(+) plasmid and transformed into *E. coli* BL21 pLysY cells, which suppress basal T7 RNA polymerase activity and tolerate the expression of potentially cytotoxic proteins^32,33^. BL21 pLysY cells containing our plasmid were grown in lysogeny broth at 37 °C until reaching their exponential phase before dilution and addition of IPTG to induce expression. Simultaneous expression of Htt and mHtt reporters within the same cellular environment was then measured using a multimode plate reader to enable direct comparative analysis under identical growth, induction, and drug conditions. Fluorescence measurements following induction were used as a quantitative proxy for expression of corresponding Htt proteins (Fig. 1c).

**Fig 1.**
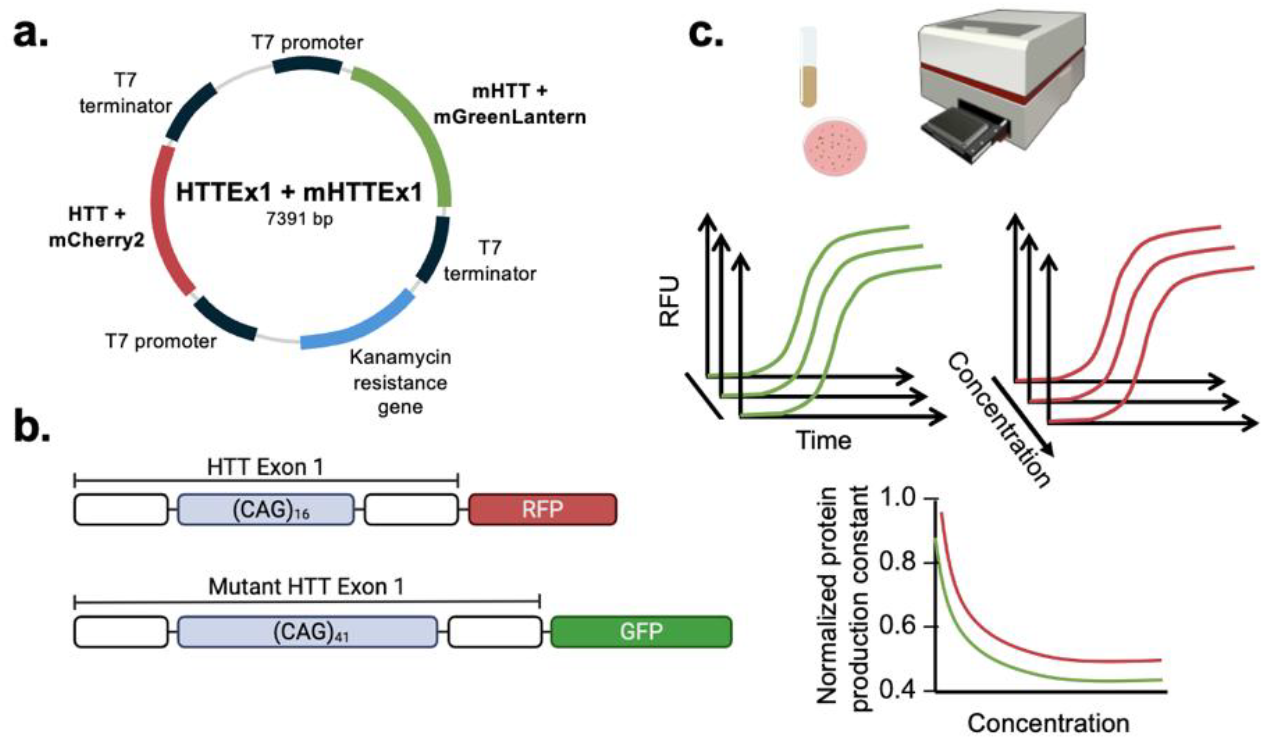
Schematic. overview of dual-reporter system with simultaneous monitoring of Htt and mHtt expression. (a) Dual-expression plasmid containing wildtype and mutant huntingtin exon 1genes, each with its own promoter and fused to a C-terminal spectrally distinct fluorescent protein, was cloned into a pET28a(+) vector with kanamycin resistance. (b) Schematic of HTT exon 1of wildtype (CAG_16_) (top) and mutant (CAG_41_) (bottom) gene with corresponding fluorescent proteins, mCherry2 and mGreenLantern, respectively. (c) Assay workflow including culturing BL21pLysY cells, plating in a 96-well plate, reading fluorescence output in a multi-mode plate reader, and subsequent fitting for quantitative analysis.

### Dose-dependent reporter responses to reference compounds

To evaluate whether our assay could detect small molecule modulation of Htt and mHtt reporter expression, two previously reported CAG repeat binding compounds, furamidine and myricetin, were tested in the dual-reporter assay. Fluorescence signals corresponding to mGreenLantern (excitation: 500 nm; emission: 521 nm)^30^ and mCherry2 (ex: 587 nm; em: 610 nm)^31^ were monitored following IPTG induction to assess compound-dependent changes in reporter fluorescence (and thus, expression of mHtt or Htt). Vehicle control wells contained 1% DMSO (matching the final DMSO concentration in compound-treated wells) to normalize the fluorescence of each reporter within the same plate (Fig. 2).

**Fig 2.**
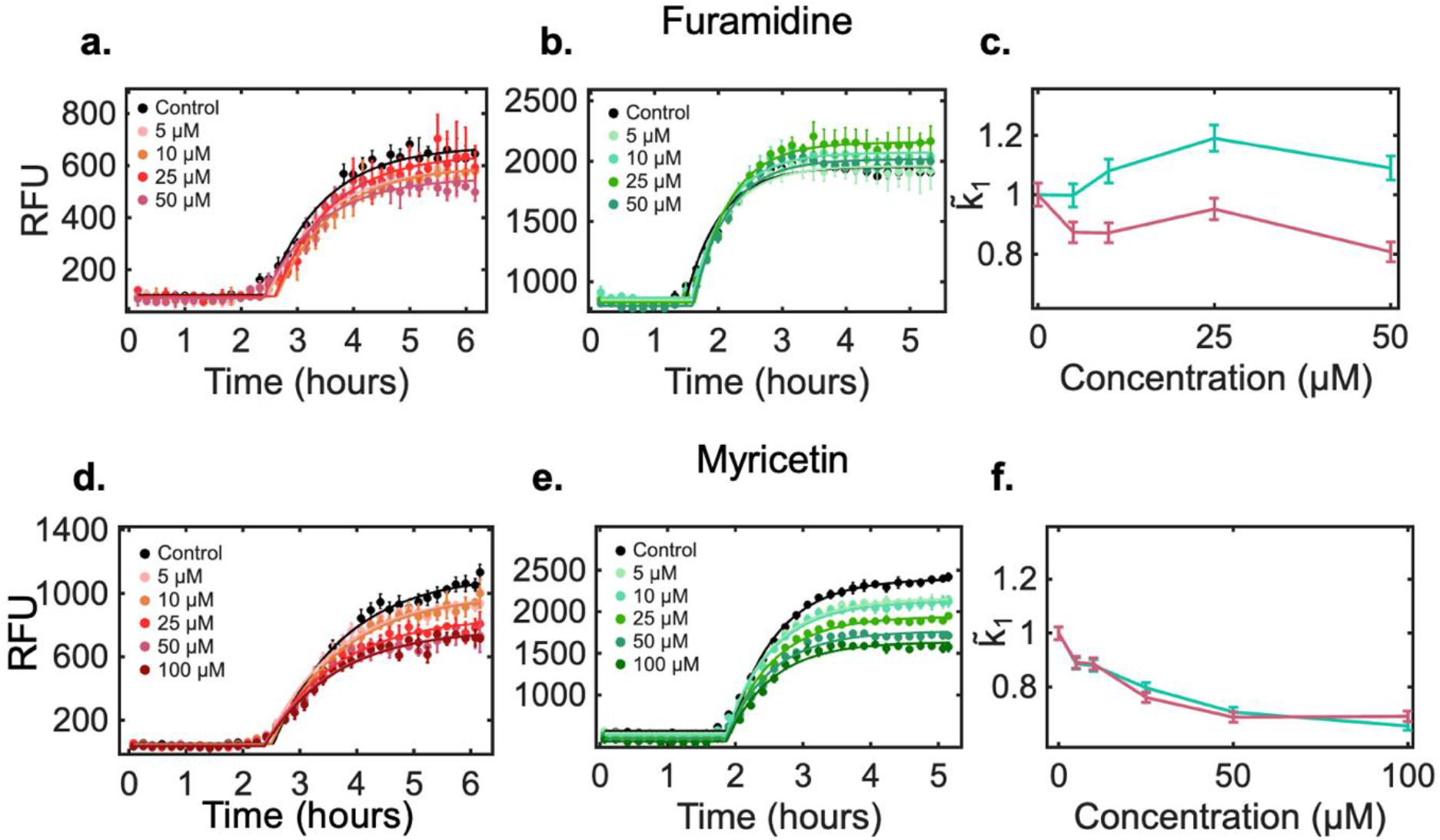
Fluorescence readings of each reporter capture expression of Htt and mHtt proteins. Time-course fluorescence signals of (a,d) mCherry2 corresponding to Htt and (b,e) mGreenLantern corresponding to mHtt in the presence of DMSO (black) or varying concentrations of (a,b) furamidine or (d,e) myricetin. Normalized 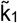 plots as a function of compound concentration for (c) furamidine and (f) myricetin for Htt (magenta) and mHtt (cyan) expression. Error bars of fluorescence data represent the standard deviation of each replicate fit (*n* = 3). Error bars for 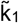 plots denote the normalized standard error of the non-linear regression fit.

Treatment with furamidine produced distinct readuts (Fig. 2a-c), decreasing the expression of the Htt reporter in a dose-dependent manner (quantified as 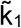 — see **Quantification analysis of reporter expression kinetics**). Expression of the mHtt reporter remained relatively unaffected relative to DMSO controls, a behavior consistent with furamidine exhibiting a higher affinity for shorter CAG repeat RNAs than longer CAG repeat RNAs^27^. In contrast, treatment with myricetin resulted in a similar dose-dependent decrease in fluorescence for both Htt and mHtt reporters without allele selectivity (Fig. 2d-f). To ensure that changes in reporter signal reflected effects on translation rather than decreased cell count, myricetin and furamidine were assessed in colony-forming unit (CFU) assays (Fig. S3). Furamidine at 100 µM reduced viable counts by 0.93 log_10_ CFU/mL relative to the vehicle, so analysis was restricted to concentrations up to 50 µM which did not significantly affect the viable cell count.

As a control against fluorophore-dependent activity, we tested furamidine in our swapped-reporter assay. Treatment with furamidine produced similar preferential decreases in Htt expression for both the original and swapped-reporter control plasmid, indicating that the observed decrease in Htt expression is associated with the Htt sequence itself rather than the fluorescent protein (Fig. S4). Since myricetin had no selectivity for one *HTT* sequence over the other, our swapped reporter system could not be used to differentiate *HTT*-specific effects from fluorescent protein-specific effects. Instead, we constructed fluorophore-only constructs without *HTT* or *mHTT* sequences to test this. Upon myricetin treatment, cells expressing either mCherry2 or mGreenLantern in the absence of either *HTT* sequence were found to have decreased fluorescence (100 µM myricetin: 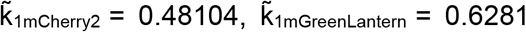, (Fig. S5). These findings indicate that furamidine exhibits *HTT*-specific effects on protein expression while myricetin ‘s observed decrease in reporter signal cannot be attributed solely to interactions with *HTT* sequences. Specifically, myricetin may induce reporter-specific effects contributing to the measured decrease in fluorescence. Therefore, it is imperative to include fluorophore controls to rule out fluorophore-induced artifacts in fluorescence-based screening assays.

### Quantification analysis of reporter expression kinetics

Each fluorescence time course was fit to a kinetic model representative of protein abundance response over time (*N*_*p*_ (*t*))^34^. The model assumes an initial lag phase before reaching a steady state of protein production and degradation, with fluorescence intensity proportional to the number of proteins (*I* (*t*) ∝ *N*_*p*_ (*t*)):

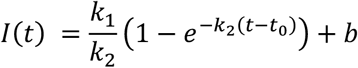

where *k*_1_is the protein production constant, *k*_2_ is the protein degradation rate constant, *t*_0_ represents the onset time at which protein production becomes exponential following the lag phase and fluorescent protein maturation, and *b* accounts for background fluorescence. For each treatment, the parameters (*k*_1_*t*_0_, and *b*) were fit independently for each small molecule condition with *k*_1_ normalized to the corresponding vehicle control to account for variability in expression and fluorescence intensities 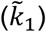. As *k*_2_ fits showed little variability across compound concentrations, they were fit globally for each compound and gene to allow for direct comparison of compound-specific effects on protein production (Table S2). Model fits are shown for furamidine and myricetin treatments in Fig. 2a-b and 2c-d, respectively. The extracted and normalized protein production constants, 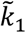 (Table 1), were subsequently plotted as a function of compound concentration (Fig 2c,f) to enable direct, quantitative analysis of dose-dependent effects on reporter expression (*n* = 3). Furamidine ‘s 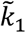 plot reflects the selective decrease in expression for the Htt reporter compared to the mHtt reporter at all concentrations examined. This supports prior *in vitro* and *in vivo* experiments in which furamidine had a higher affinity for *HTT* over *mHTT*^27^ but also crucially establishes that in the presence of both sequences, furamidine preferentially targets *HTT*. As expected, myricetin ‘s 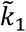 plot shows a dose-dependent but nonspecific decrease for both mHtt and Htt reporters (Fig. 2f).

**Table 1.**
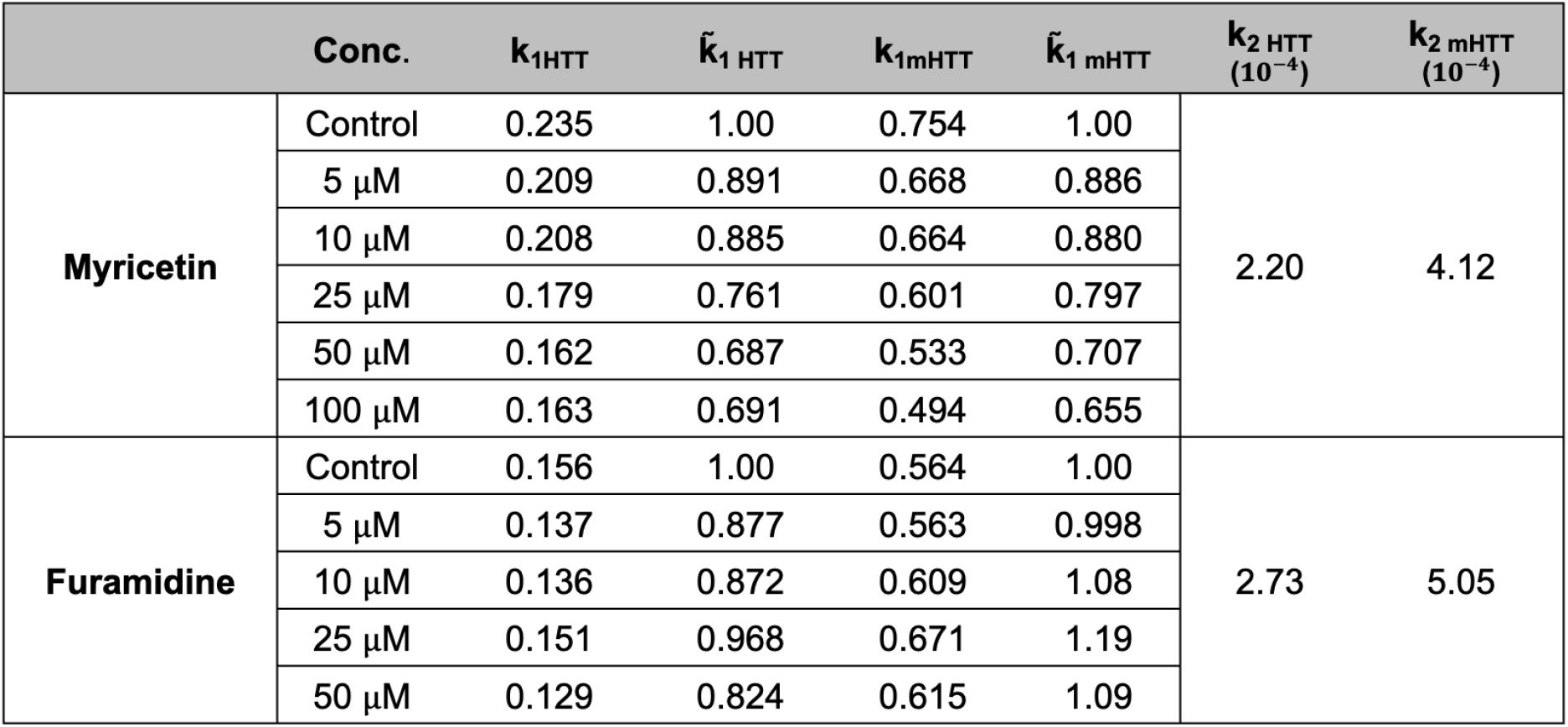
Fitted parameters of raw (k_1_) and normalized 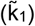 protein production constants for Htt and mHtt expression with 5, 10, 25, 50, or 100 *μ*M of myricetin and furamidine. Protein degradation rate constant (k_2_) for Htt and mHtt expression for myricetin (top) and furamidine (bottom).

To confirm assay sensitivity, we used 50 µg/mL spectinomycin, a bacteriostatic antibiotic that binds the 30S ribosomal subunit and blocks the translocation step of translation. Spectinomycin inhibited the increase in reporter fluorescence compared to the vehicle controls (Fig. 3). Colony-forming unit assays were performed and spectinomycin was found to reduce viable counts by 0.8 log_10_ CFU/mL relative to the vehicle control (Fig. S6)^35^, but below the 3log_10_ threshold for bactericidal activity^36^. Consequently, spectinomycin serves as a reference for maximal inhibition of reporter expression, validating our assay ‘s sensitivity and capability at detecting translation inhibitors.

**Fig 3.**
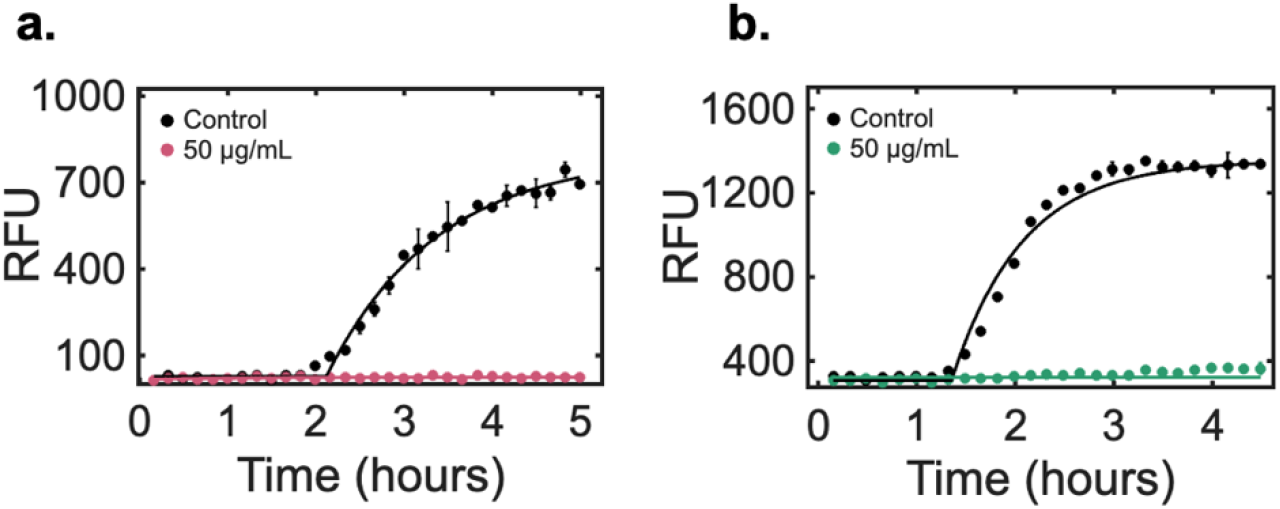
Time-course fluorescence signals of (a) mCherry2 corresponding to Htt and (b) mGreenLantern corresponding to mHtt in the presence of DMSO (black) or 50 μg/mL spectinomycin. Error bars represent the standard deviation of replicates (*n* = 3).

### Conformational changes of RNA structure in the presence of furamidine or myricetin

To determine whether furamidine and myricetin directly interact with CAG repeat RNA prior to translation, we used circular dichroism (CD) spectroscopy thermal melts to track changes in RNA unfolding in the presence of furamidine or myricetin compared to the DMSO control^20,24,37^. Thus, by measuring the CD signal at 260 nm (corresponding to folded RNA) across a temperature range of 25°C – 95°C, changes in RNA stability were assessed via the apparent melting temperature (T_m_), defined as the temperature in which half of the RNA secondary structure has dissociated (Fig. 4). A compound-induced change in T_m_ would provide biophysical evidence of direct RNA interaction and support the hypothesis that destabilization of CAG repeat transcripts underlies the decrease in Htt expression seen in the dual-expression cellular assay.

**Fig 4.**
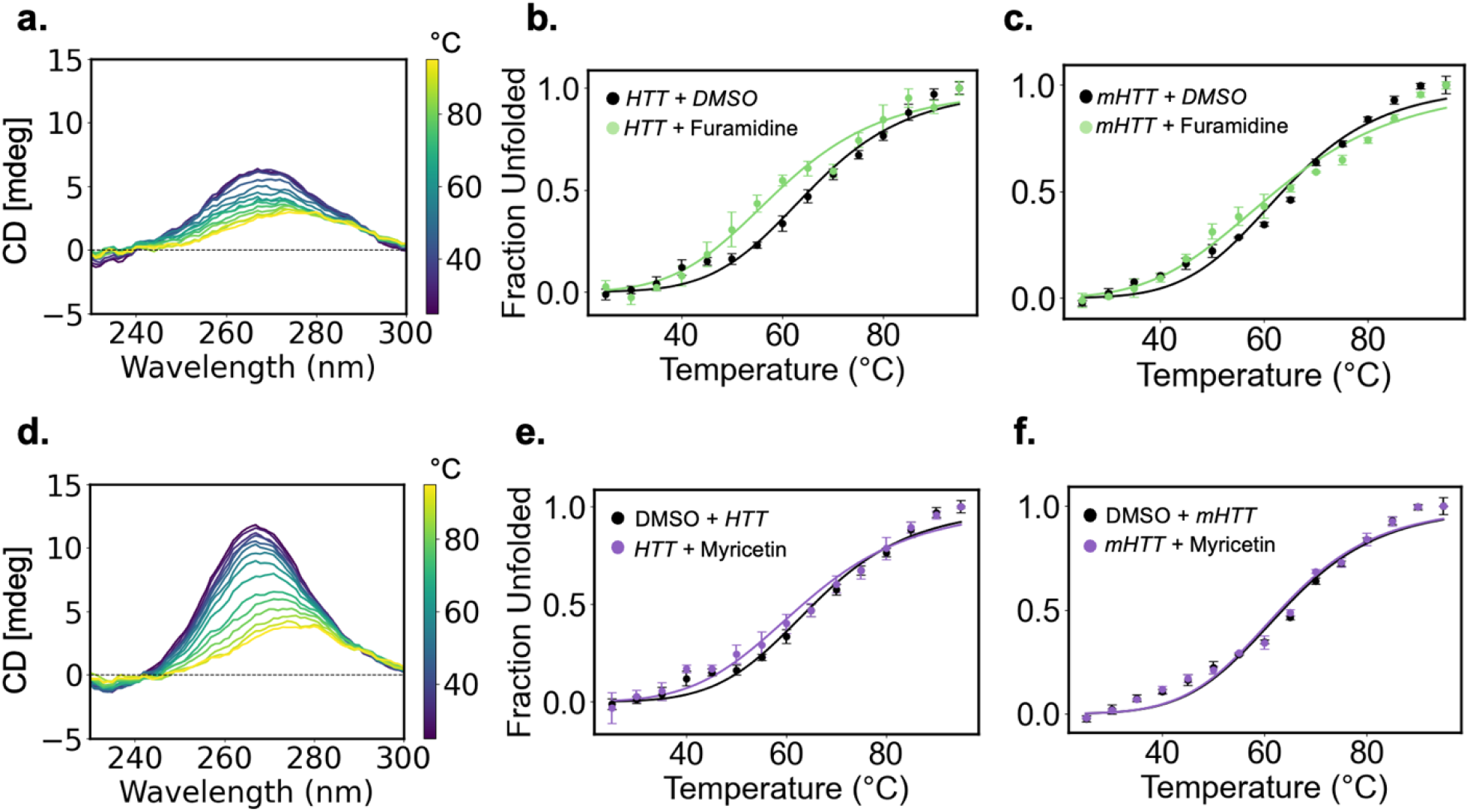
Biophysical assessment of compound-induced changes in thermal stability of expanded CAG repeat transcripts. Representative CD spectra of (a) *HTT* RNA thermal melt in the presence of furamidine and (d) *mHTT* in the presence of myricetin. Thermal denaturation curve of (b) *HTT* RNA in the presence of furamidine (green), DMSO (control, black) and (e) myricetin (purple). Thermal denaturation curve of (c) *mHTT* RNA in the presence of furamidine, DMSO or (f) myricetin. Error bars represent the average of replicates (*n* = 3). Spectra are averaged from three technical replicates.

Relative to the DMSO control, furamidine produced distinct effects on the thermal stability of *HTT* RNA, decreasing the T_m_ by ∼ 6 °C (Fig. 4b). In contrast, the *mHTT* RNA exhibited little to no change in thermal stability upon furamidine treatment (*HTT*: T_m_ = 59.4 ±0.9 °C; *mHTT*: T_m_ = 65.5 ±0.9 °C, relative to DMSO controls of T_m_= 65.5 ±0.9 °C and 64 ±1°C, respectively) (Fig. 4c). These data demonstrate that furamidine has a stronger effect on RNA stability of *HTT* over *mHTT*, suggesting that this *HTT* destabilization leads to a disruption in translation and consistent with a selective decrease in Htt protein expression observed in our dual-expression assay. In contrast, myricetin produced no significant effects on RNA thermal stability for either *HTT* or *mHTT* (*HTT*: T_m_ = 63 ±1°C; *mHTT*: T_m_ = 63.1±0.9 °C, relative to DMSO controls of T_m_= 65.5 ±0.9 °C and 64 ±1°C, respectively) under the conditions examined (Fig. 4e,f). The lack of a compound induced change in T_m_ indicates that myricetin does not directly interact with *HTT* or *mHTT* RNA, suggesting that its effect on the reporter fluorescence in the dual-expression assay is attributable to fluorescent-protein specific interactions rather than RNA targeting.

Together, these results demonstrate that CD spectroscopy can distinguish compound-specific effects on the stability of *HTT* and *mHTT* RNA, with furamidine differentially destabilizing the *HTT* RNA and myricetin leaving both transcripts unaffected. Although the similar T_m_ values observed *HTT* and *mHTT* transcripts under DMSO control conditions may reflect the dilute, uncrowded nature of our measurements, the preferential destabilization of *HTT* RNA in the presence of furamidine is consistent with its selective downregulation of Htt expression in the dual-expression assay. For myricetin, the absence of any RNA perturbation by CD indicates that its effect cannot be attributed to RNA interactions and instead points to a fluorophore-specific interaction.

### Cell-free protein synthesis system distinguishes cell-independent compound-specific effects on HTT reporter expression

Although CD experiments indicate furamidine directly interacts with CAG repeat RNA, we sought to determine whether furamidine ‘s effect on protein expression resulted from a direct interaction with CAG repeat RNA as opposed to an interaction with other cellular processes in the protein production. To isolate the translation of transcripts, we evaluated our small molecules in a cell-free protein synthesis system (CFPS) using purified *mHTT* and *HTT* transcripts. *In vitro* translation included simultaneous expression of Htt and mHtt (fused to their spectrally distinct fluorophores) at 0.25 µM (0.5 µM total) in the presence of 50 µM of compounds, or equivalent DMSO control, replicating drug:RNA ratios used in CD experiments. As with the cellular assay, reporter fluorescence was measured over time to monitor protein expression, and 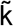 parameters were extracted (Fig. 5). Strikingly, furamidine exhibited a preferential decrease in Htt reporter expression compared to mHTT 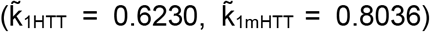 (Fig 5a-b), validating our hypothesis that the decrease in Htt reporter expression in the presence of furamidine is due to an effect on translation in our dual-expression assay. In contrast, myricetin demonstrated differential effects in the cell-free system, seemingly decreasing the translation of the mHtt reporter more than the Htt reporter 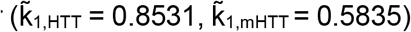 (Fig. 5c-d). These inconsistencies between the dual-expression cellular assay, CD data, and CFPS results further support evidence that myricetin influences the fluorescent protein reporters themselves.

**Fig 5.**
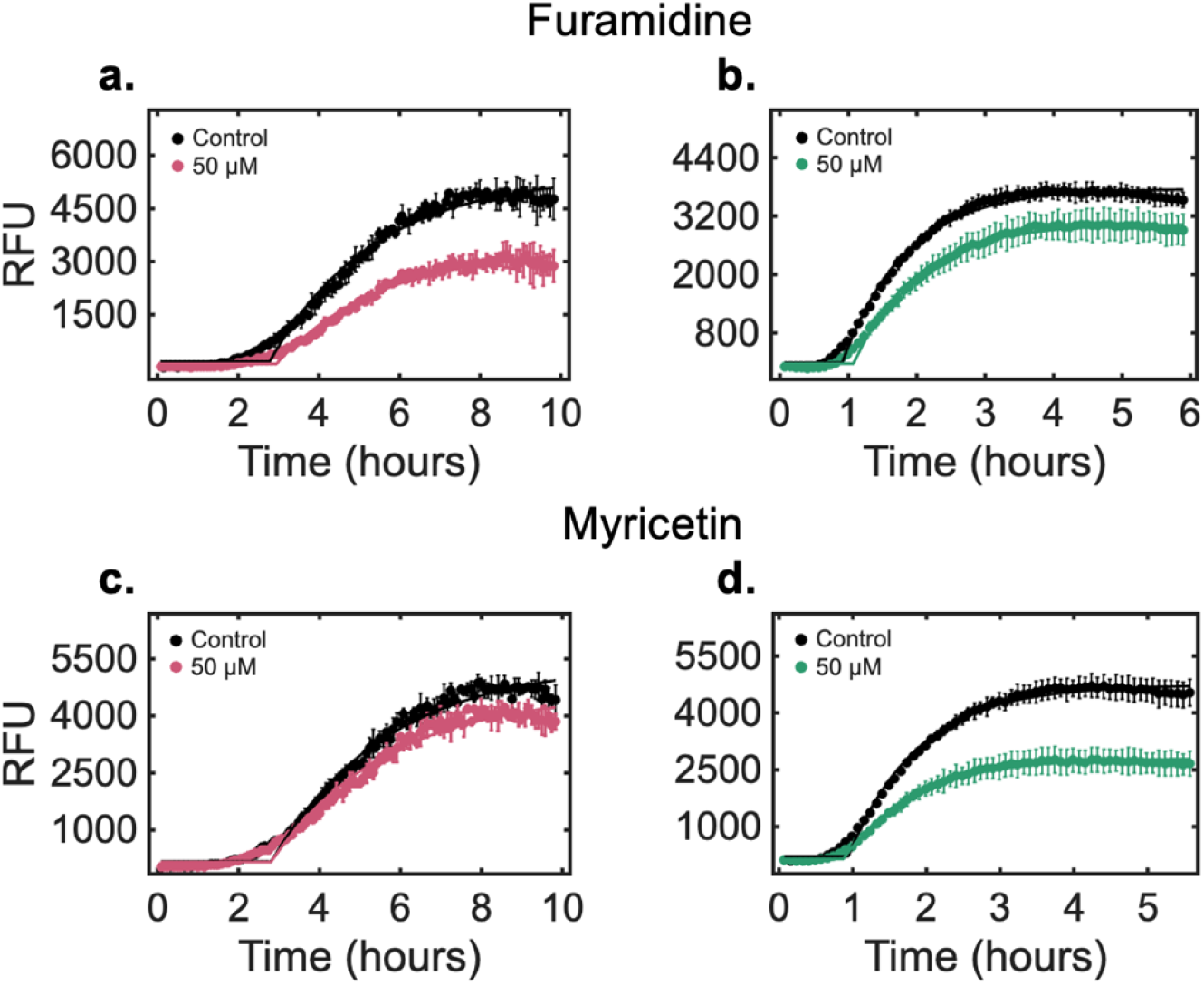
Cell-free protein synthesis reveals that the effect of furamidine is primarily due to translation. Time course fluorescence signals of (a,c) mCherry2 corresponding to Htt and (b,d) mGreenLantern corresponding to mHtt in the presence of DMSO (black) or 50 *μ*M of (a,b) furamidine or (c,d) myricetin. Error bars represent the standard deviation of replicates (*n* = 3).

Collectively, these experiments demonstrate that the dual expression assay, complemented by CD spectroscopy and CFPS, can distinguish genuine modulators of Htt expression from fluorophore or compound-specific artifacts. Furamidine ‘s consistent allele selectivity for *HTT* across all three experiments confirms that our workflow correctly identifies sequence and structural-dependent translational effects. The limitations of our assay are evident in that fact that some small molecules (myricetin) may interact with the fluorescent reporters themselves, necessitating the complementary workflow to accurately attribute changes in reporter fluorescence to specific mechanisms. Although our workflow has only been benchmarked against furamidine and myricetin, it can be readily adapted to screen existing or proposed small molecules.

## Discussion and Conclusion

Small molecule modulators of RNA present an attractive therapeutic alternative due to improved tissue bioavailability and cellular permeability. By exploiting the highly stable RNA hairpins formed by expanded CAG repeats, relative to the less stable, shorter, non-pathogenic tracts, small molecules can preferentially target disease-associated RNA transcripts. However, discovery of potential small molecule therapeutics for HD requires effective drug screening in a high-throughput, reproducible system capable of detecting both compound efficacy and transcript selectivity. Conventional workflows relying on *in silico* modeling, surface plasmon resonance, isothermal titration calorimetry, FRAP, or fluorescence titration assays are often labor-intensive, technically demanding, and may fail to measure the translation inhibition in a high-throughput manner, limiting the biological relevance of these assays. For this purpose, we have developed and validated a dual-expression fluorescence reporter platform capable of simultaneously monitoring wildtype and mutant Htt reporter expression in a bacterial system. This platform provides a rapid and quantitative strategy for evaluating small molecules ‘ ability to modulate Htt expression while directly comparing Htt and mHtt expression under identical experimental conditions.

The development of small molecule therapeutics targeting pathogenic RNA structures offers a promising strategy in mitigating neurodegenerative pathology while overcoming delivery issues present in current biological therapeutics. Because several trinucleotide repeat expansion disorders (TREDs) have expanded RNA sequences that fold into distinct secondary structures, this strategy has broad therapeutic applicability. For example, neurological diseases such as fragile X-associated tremor ataxia syndrome are triggered by CGG repeats that can adopt stable sequence-dependent RNA structures for preferential targeting by small molecules^2,26^. Similarly, spinocerebellar ataxia type 3 and Huntington ‘s disease (HD) stem from expanded CAG repeats^24,38^, suggesting that molecules developed through this approach may serve as therapeutics across both disorders.

A notable limitation of this assay is the use of the bacterial system. Although it is efficient in isolating a drug ‘s effect on Htt and mHtt production, it cannot fully recapitulate transcription and translation as it occurs physiologically in neuronal cells; therefore, small molecules found to decrease Htt translation in the bacterial system may not have the same effect in neuronal cells. Nevertheless, our bacterial system is a sufficient initial screen as a biologically relevant, structural RNA assay that can be utilized as a starting point for more complex cellular or animal models in the future.

Future work in optimizing small molecule candidates is needed to increase selectivity for expanded CAG hairpins and minimize off-target interactions with wildtype transcripts. To this end, *in silico* screening methods can be used to model the compounds known to cross the blood-brain barrier and bind to pathogenic RNA. Additionally, translating this platform to a mammalian context would give greater biological relevance to confirm a candidate compound ‘s efficacy and long-term effects on proteotoxicity. Finally, since pathogenic trinucleotide repeat expansions are implicated in many neurodegenerative disorders, our high-throughput screening platform may be applicable across other repeat-expansion disorders such as myotonic dystrophy and spinocerebellar ataxias.

## Materials and Methods

### Compounds

Myricetin and furamidine were purchased from Millipore Sigma (St. Louis, Missouri).

### Plasmids and cloning

Dual-expression HTTQ16/HTTQ41 plasmid constructs were designed and purchased from GenScript (Piscataway, New Jersey). Fluorophore-only control constructs were cloned using restriction enzyme digest and ligation using the XhoI enzyme from New England Biolabs (Ipswich, Massachusetts) into a pET28a(+) plasmid.

### Bacterial cells

BL21 pLysY E. coli cells were purchased from New England Biolabs (Ipswich, Massachusetts).

### Dual-expression bacterial cell plate reader assay

Colonies were picked from overnight incubation to inoculate 5 mL cultures of lysogeny broth containing 50 µg/mL kanamycin. Cultures were grown to an OD_600_ of 0.6 and diluted to 0.1 before addition of 1mM IPTG. Using a multichannel pipette, 198 µL of culture was aliquoted into each well of a 96-microwell plate. Either 2 µL of DMSO for the control data or 2 µL of compound at varying concentrations was added to each well in triplicate. Phosphate-buffered saline (PBS) was pipetted into the spaces between wells to ensure equilibrated heating upon incubation. A multi-mode plate reader was used (BioTek Neo2 Agilent) and set at 37 °C with orbital shaking (frequency: 237 cpm (4mm)). Fluorescence filters were set at excitation: 500/10; emission: 521/10 for mGreenLantern and excitation: 587/10; emission: 610/10 for mCherry2 with the gain set to 100 and 130, respectively. Fluorescence intensity measurements were taken at 10-minute intervals while cultures were grown for 6-8 hours. Replicate-averaged time-intensity curves were obtained for quantification.

### Kinetic model fitting

The kinetic model is based upon a differential equation that captures protein translation:

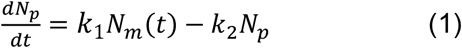

where *N*_*p*_ corresponds to the number of a particular protein, *k*_1_is the mRNA-dependent protein expression constant, *N*_*m*_ (*t*) is the number of mRNA transcripts for a particular protein, and *k*_2_ is the protein degradation rate constant.^34^ If we reasonably assume *N*_*m*_ (*t*) rapidly reaches steady state with a strong inducer and fluorescence intensity is directly proportional to the number of proteins (i.e. *I* (*t*) ∝ *N*_*p*_ (*t*) with sufficiently rapid folding times relative to the length of the assay) we can solve Eq. 2:

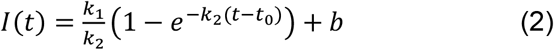

where *k*1is the new protein expression rate constant, *b* corresponds to an arbitrary background constant and *t*_0_ is the time in which protein production becomes exponential. While Eq. 2 takes into account several simplifying assumptions, it sufficiently captures the production constant and degradation rate constant of both mHTT and HTT in an early time window immediately after induction.

### Real-time PCR

Total RNA was isolated using the Direct-Zol RNA purification kit (Zymo). cDNA was synthesized using M-MLV reverse transcriptase (Promega) with random primers (Promega), incubating for 10 min at 23°C, 60 min at 37°C, and 5 min at 70°C. Real-time PCR was performed using the EvaGreen PCR master mix (Biotium). The qPCR was performed using an AriaMx Real-Time PCR system (Agilent) using the following conditions: initial denaturation of 3 min at 95°C, followed by 40 cycles of 20 s at 95°C, 20 s at 60°C, and 20 s at 72°C for annealing and extension. Subsequent dissociation curves were run at 95°C for 15 s, 60 °C for 15 s and 95°C for 15 s. Samples were analyzed in triplicate and were normalized to the reference gene idnT (L-idonate/5-ketogluconate/gluconate transporter) endogenously expressed in E. coli^39^. The following primers were used: idnT-forward, CTGTTTAGCGAAGAGGAGATGC, idnT-reverse, ACAAACGGCGGCGATAGC; HTT-mch2 forward, agttcatgcgcttcaaggtg, HTT-mch2 reverse, aagtagtcggggatgtcg; mHTT-mGL forward, aagaacgcaccatcagctt, mHTT-mGL reverse, cgttcttctgtttgtccgca.

### Colony-forming unit assay

Cell cultures at an OD_600_ of 0.6 were diluted to 0.1 with an IPTG concentration of 1 mM added. Either DMSO (control) or drug was included and the cultures were allowed to grow for 4-6 hours in a shaking incubator at 37 °C. Serial dilutions in PBS were performed from 100 µL culture (1×10−^1^ to 1×10−^6^ ). The last three dilutions were plated on agar plates containing 50 µg/mL kanamycin in duplicate. Colonies were left to grow at 37°C overnight and enumerated after 18-20 hours of growth. Colony numbers were multiplied by dilution factor to calculate colony-forming unit log_10_(CFU)/mL.

### *In vitro* transcription and RNA purification

*In vitro* transcription was performed using the New England Biolabs HiScribe T7 High Yield RNA Synthesis kit as commercially described. For the DNA templates, separate plasmids containing HTT-mCherry2 or mHTT-mGreenLantern were linearized using restriction enzyme digest with XhoI. DNase treatment of the subsequent IVT product was performed before spin-column purification using the New England Biolabs Monarch Spin RNA cleanup kit according to manufacturer ‘s instructions.

### Cell-free protein synthesis

Cell-free protein synthesis of RNA templates was performed using the New England Biolabs NEBExpress Cell-free *E. coli* Protein Synthesis System according to manufacturer ‘s instructions. Reactions were performed in a 96 well half area plate in a multi-mode plate reader (BioTek Neo2 Agilent) to monitor expression of reporter protein with identical conditions to the dual-expression bacterial assay, adjusting the gain as needed.

### Circular dichroism

Circular dichroism (CD) experiments were performed on a Jasco J-1100 spectrometer (Jasco, Inc. Mary ‘s Court, Easton, Maryland, US) equipped with a Peltier temperature controller at successively increasing temperatures of 1°C increments from 25 °C to 95 °C, collecting spectra every 5 °C. The purified RNA transcripts of HTT exon 1 contained either 16 or 41 CAG repeats corresponding to *HTT* or *mHTT*, respectively. RNA concentration was held constant at 0.75 µM in 10 mM sodium phosphate pH 7.2, 100 mM potassium chloride, and 5 mM magnesium chloride with 75 µM drug or DMSO vehicle control. All CD experiments were performed in a quartz cuvette with a 0.1 cm pathlength, and spectra were recorded at 1 nm intervals from 210 nm to 300 nm with a 1 nm – slit width and averaged over three scans. Data were normalized by subtracting the buffer + compound/DMSO CD spectra from RNA CD spectra. Thermal denaturation curves were plotted using the averaged ellipticity signal at 260 nm, assuming complete higher-order structure formation at 25 °C and complete higher-order structure denaturation at 95 °C. By fitting data to the Hill equation (Eq. 3), the denaturation midpoint (T_m_) which can be used as a measure of thermal stability, was obtained where n represents the Hill coefficient.

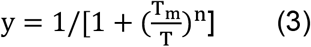

