## Supplementary Information file for "Dual-reporter assay for selective small molecule inhibition of expanded toxic huntingtin"

*Corresponding author

### Supplementary Information

| | |
|---|---|
| Myricetin<br>MW: 318.24 g/mol | 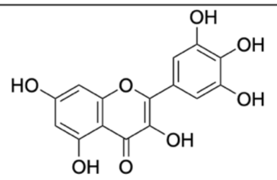 |
| Furamidine<br>MW: 377.27 g/mol | 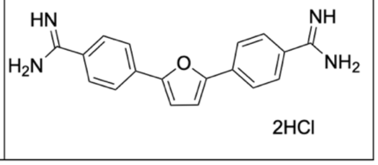 |

**Supplementary Table S1** Myricetin and furamidine molecular weights and chemical structures.

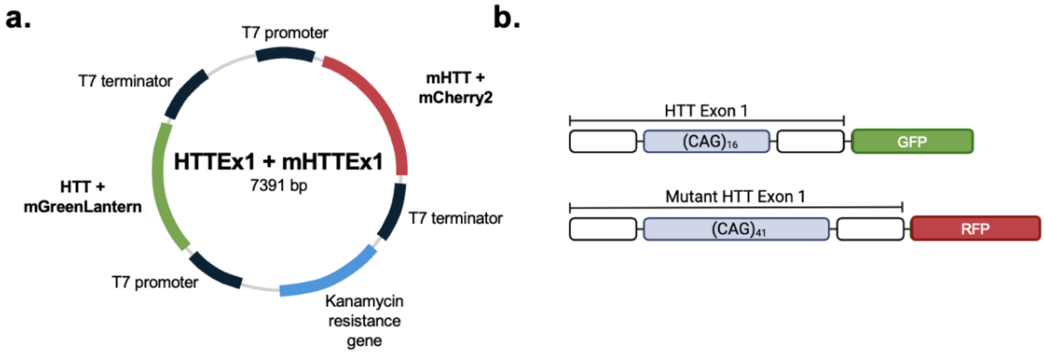


**Supplementary Figure S1 Schematic overview of swapped dual-reporter system with simultaneous monitoring of HTT and mHTT expression**. (a) Dual-expression plasmid containing wildtype and mutant huntingtin exon 1 genes with kanamycin resistance in a pET28a(+) vector. (b) Schematic of HTT exon 1 of wildtype ($CAG_{16}$) (top) and mutant ($CAG_{41}$) (bottom) gene with corresponding fluorescent proteins, mGreenLantern and mCherry2, respectively.

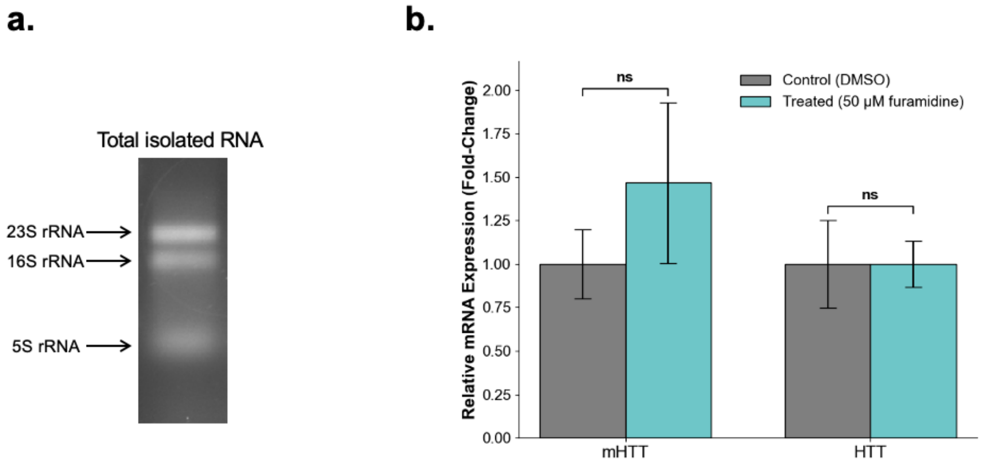


**Supplementary Figure S2.** (a) Total isolated RNA from BL21 pLysY cells expressing dual-expression plasmid in the presence of 50 $\mu$M furamidine with characteristic 23s, 16s, and 5s rRNA bands. (b) RT-qPCR for quantification of *mHTT* and *HTT* transcripts using relative quantification with idnT gene.

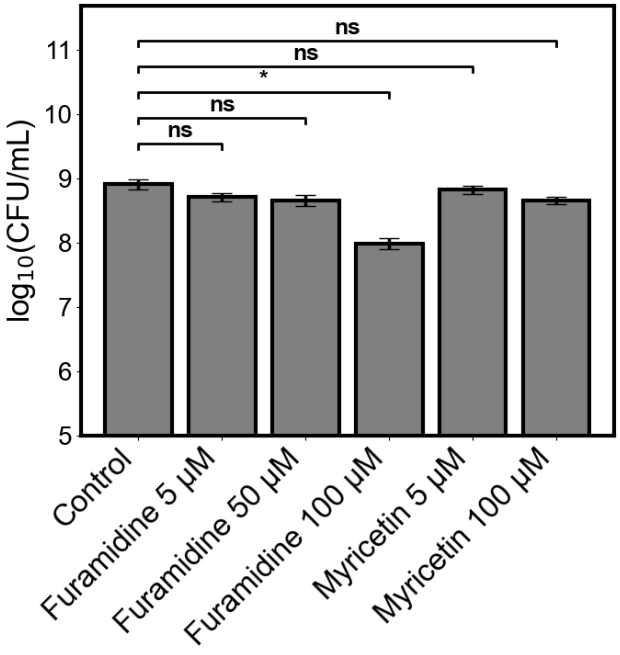


**Supplementary Figure S3** Colony-forming unit assay for viable cell counts after furamidine or myricetin treatment. Control data corresponds to cells receiving the vehicle alone (DMSO). At 100 $\mu$M, furamidine decreased viable counts significantly by 0.93 $log_{10}$ CFU/mL. For this reason, 100 $\mu$M furamidine treatment was omitted from analysis.

| | Conc. | $k_{1HTT}$ | $\tilde{k}_{1\ HTT}$ | $k_{1mHTT}$ | $\tilde{k}_{1\ mHTT}$ | $k_{2HTT}$ $(10^{-4})$ | $k_{2mHTT}$ $(10^{-4})$ |
|---|---|---|---|---|---|---|---|
| **Myricetin** | Control | 0.211 | 1.00 | 0.886 | 1.00 | 1.88 | 3.89 |
| | 5 µM | 0.215 | 1.02 | 0.648 | 0.902 | 2.28 | 3.98 |
| | 10 µM | 0.183 | 0.868 | 0.645 | 0.898 | 1.84 | 3.99 |
| | 25 µM | 0.189 | 0.898 | 0.597 | 0.831 | 2.37 | 4.09 |
| | 50 µM | 0.183 | 0.870 | 0.666 | 0.927 | 2.60 | 5.32 |
| | 100 µM | 0.168 | 0.797 | 0.645 | 0.898 | 2.30 | 5.59 |
| **Furamidine** | Control | 0.154 | 1.00 | 0.534 | 1.00 | 2.69 | 4.76 |
| | 5 µM | 0.160 | 0.947 | 0.607 | 1.14 | 2.95 | 5.49 |
| | 10 µM | 0.134 | 0.865 | 0.634 | 1.19 | 2.67 | 5.29 |
| | 25 µM | 0.129 | 0.835 | 0.608 | 1.14 | 2.25 | 4.52 |
| | 50 µM | 0.137 | 0.890 | 0.671 | 1.3 | 2.95 | 5.57 |
| **Furamidine*** | Control | 0.114 | 1.00 | 0.553 | 1.00 | 1.84 | 5.69 |
| | 5 µM | 0.084 | 0.738 | 0.532 | 0.961 | 1.48 | 7.37 |
| | 10 µM | 0.088 | 0.772 | 0.558 | 1.01 | 1.51 | 7.77 |
| | 25 µM | 0.107 | 0.939 | 0.590 | 1.07 | 1.71 | 7.33 |
| | 50 µM | 0.095 | 0.834 | 0.606 | 1.10 | 1.82 | 8.28 |

**Supplementary Table S2.** Fitted parameters of raw ($k_1$) and normalized ($\tilde{k}_1$) protein production constants for Htt and mHtt expression with 5, 10, 25, 50, or 100 $\mu$M of myricetin or furamidine when fitting the protein degradation rate constant ($k_2$) independently for each concentration. $K_2$ values remain largely unchanged between concentrations, allowing for global fits to assess compound-specific effects on protein production.

*Denotes furamidine treatment in the swapped reporter assay

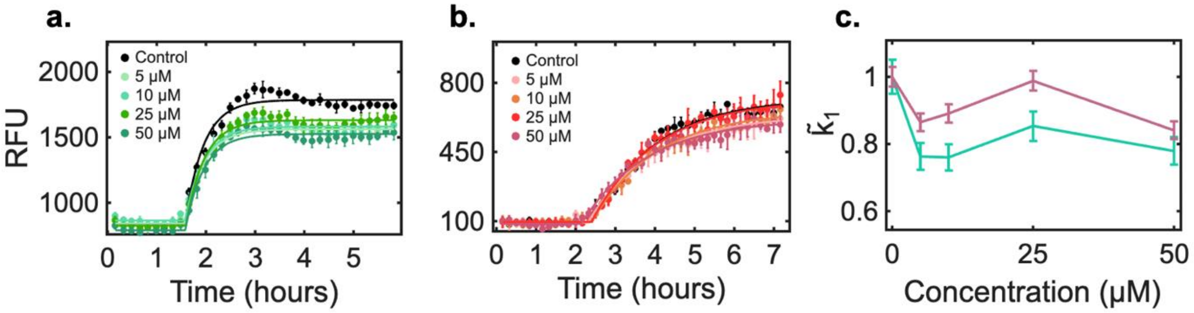


**Supplementary Figure S4**. **Fluorescence readings of the swapped-reporter system with furamidine.** Time-course fluorescence signals of (a) mGreenLantern corresponding to Htt and (b) mCherry2 corresponding to mHtt in the presence of DMSO (black) or varying concentrations of furamidine. (c) Normalized $k_1$ plots as a function of furamidine concentration. Error bars of fluorescence data represent the standard deviation of each replicate fit ($n = 3$). Error bars for $\tilde{k}_1$ plots denote the normalized standard error of the non-linear regression fit.

| | Conc. | $k_{1HTT}$ | $\tilde{k}_{1\ HTT}$ | $k_{1mHTT}$ | $\tilde{k}_{1\ mHTT}$ | $k_{2\ HTT}$ $(10^{-4})$ | $k_{2\ mHTT}$ $(10^{-4})$ |
|---|---|---|---|---|---|---|---|
| | Control | 0.105 | 1.00 | 0.701 | 1.00 | | |
| | 5 μM | 0.091 | 0.866 | 0.531 | 0.758 | | |
| **Furamidine*** | 10 μM | 0.094 | 0.893 | 0.530 | 0.756 | 1.66 | 7.36 |
| | 25 μM | 0.104 | 0.992 | 0.592 | 0.845 | | |
| | 50 μM | 0.088 | 0.841 | 0.542 | 0.773 | | |

**Supplementary Table S3**. Fitted parameters of raw ($k_1$) and normalized ($\tilde{k}_1$) protein production constants for Htt and mHtt expression with 5, 10, 25, or 50 μM furamidine in the swapped reporter system (denoted by *). Protein degradation rate constant ($k_2$) for Htt and mHtt expression fit globally for each gene

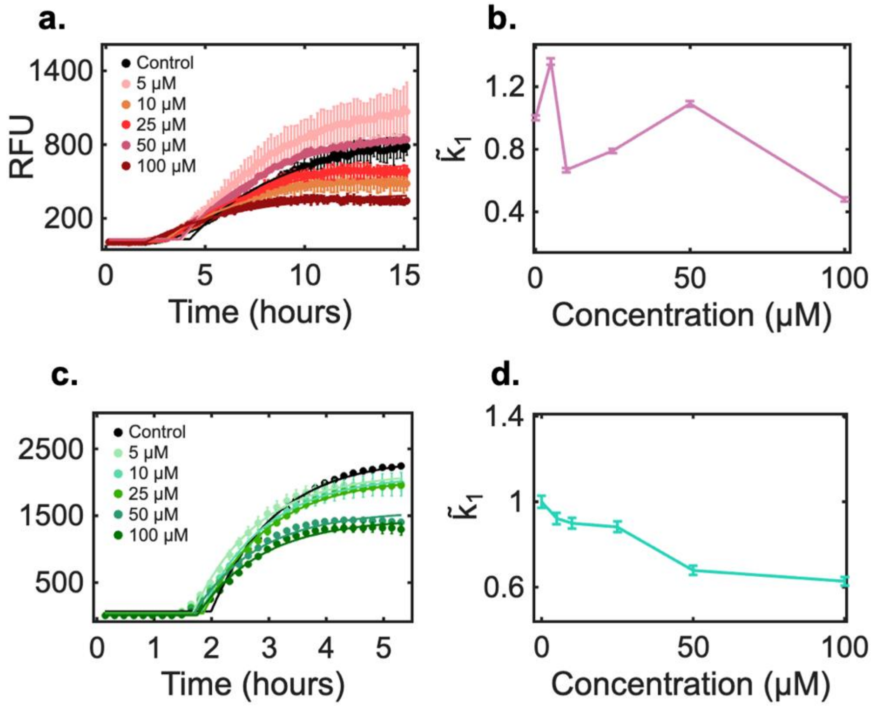


**Supplementary Figure. S5 Fluorescence signals of fluorophore-only control constructs with myricetin**. Time course fluorescence signals of (a) mCherry2 or (c) mGreenLantern in the presence of 5, 10, 25, 50, or 100 $\mu$M myricetin or DMSO control (black). Corresponding normalized $\tilde{k}_1$ plots as a function of myricetin concentration for (b) mCherry2 plots and (d) mGreenLantern plots. Error bars of fluorescence data represent the standard deviation of each replicate fit (*n* = 3). Error bars for $\tilde{k}_1$ plots denote the normalized standard error of the non-linear regression fit.

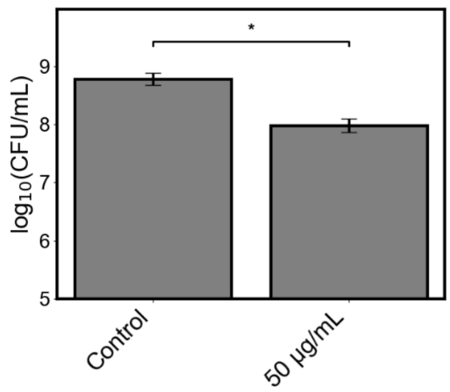


**Supplementary Figure S6** Colony-forming unit assay for viable cell counts of cells receiving 50 μg/mL spectinomycin. Control data corresponds to cells receiving the vehicle alone (DMSO).

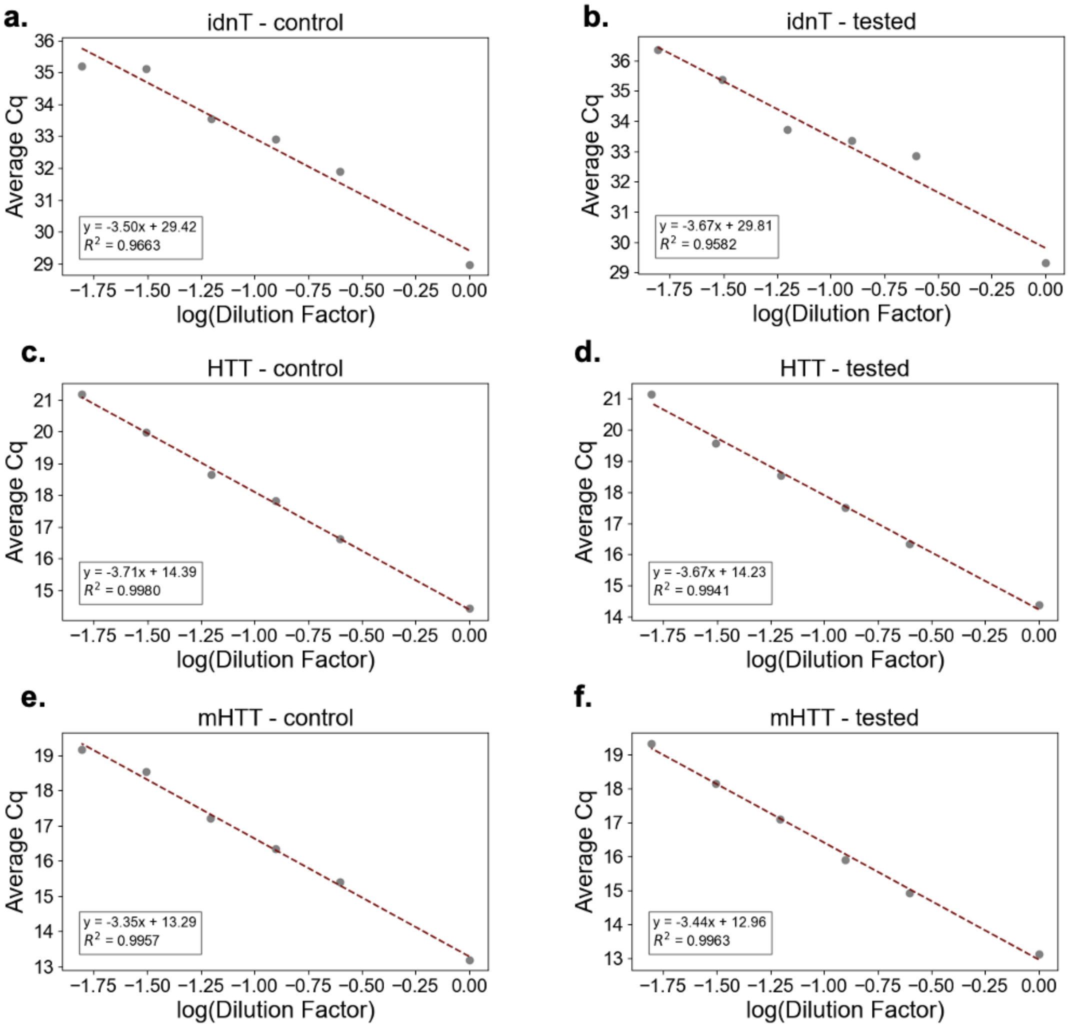

**Supplementary Figure S7.** Standard curve dilutions for primer efficiency calculations between control and tested samples. Primer sets include idnT (a,b), HTT (c,d), and mHTT (e,f). Primer efficiency was calculated using $E = \left(10^{-\left(\frac{1}{slope}\right)}\right) \times 100$.

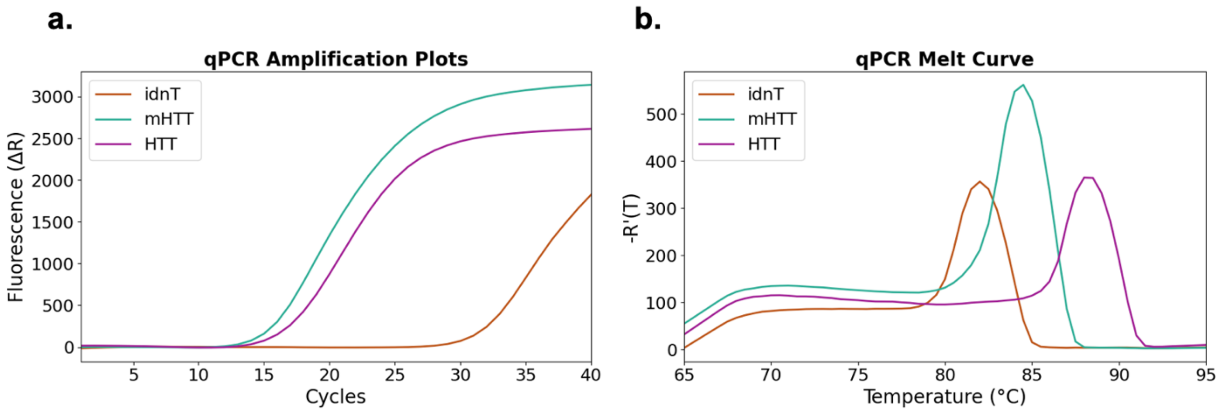


**Supplementary Figure S8.** Representative qPCR plots of amplification (a) and dissociation (b) for idnT, HTT, and mHTT confirm specific amplification of a single product.

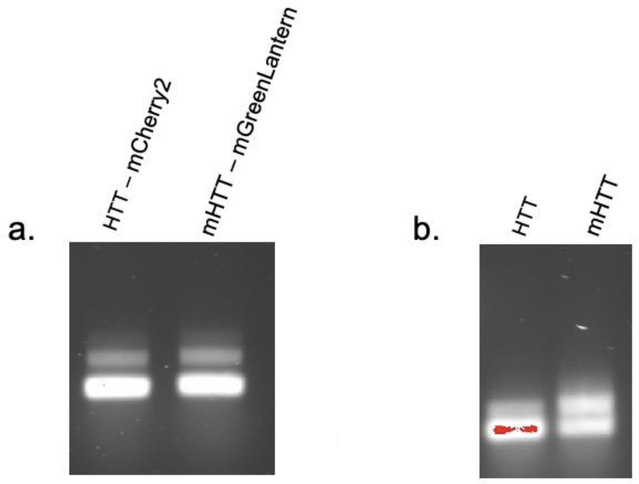


**Supplementary Figure S9.** *In vitro* transcribed RNA of *HTT – mCherry2* and *mHTT – mGreenLantern* for cell-free protein synthesis experiments (a) and *in vitro* transcribed *HTT* and *mHTT* RNA for circular dichroism experiments (b). Two bands observed for one species of RNA is assumed to be due to partially folded and unfolded RNA states present in expanded CAG-repeat RNA.

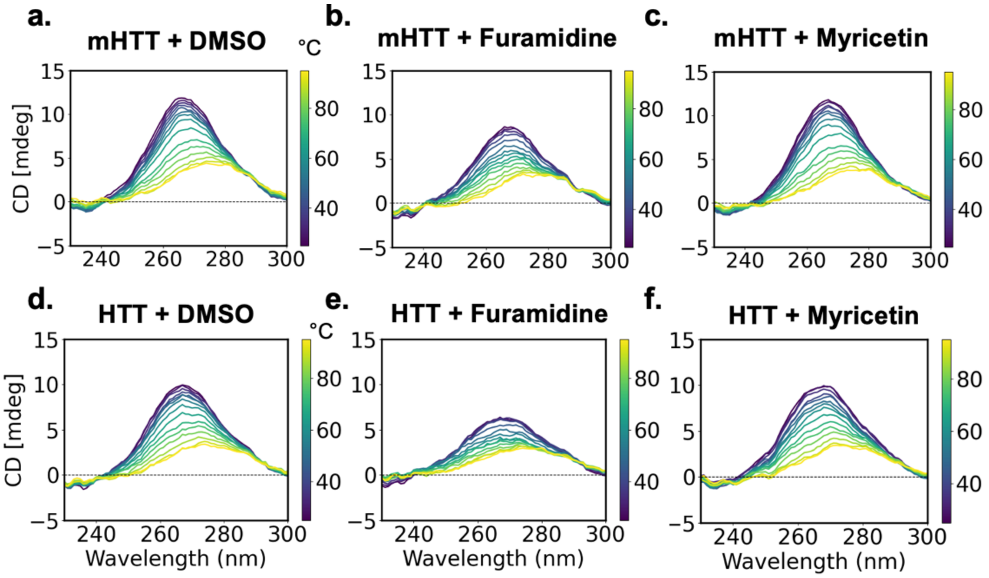


**Supplementary Figure S10.** CD spectra of *HTT* and *mHTT* RNA in the presence of either DMSO (control), furamidine, or myricetin. Spectra are averaged from three technical replicate scans.
